# The heart microbiome of insectivorous bats from Central and South Eastern Europe

**DOI:** 10.1101/2020.04.30.069815

**Authors:** Alexandra Corduneanu, Andrei Daniel Mihalca, Attila D. Sándor, Sándor Hornok, Maja Malmberg, Natalia Pin Viso, Erik Bongcam-Rudloff

**Affiliations:** Department of Parasitology and Parasitic Diseases, University of Agricultural Sciences and Veterinary Medicine of Cluj-Napoca, Romania; Department of Parasitology and Zoology, University of Veterinary Medicine, Budapest, Hungary; Section of virology, Department of Biomedical Sciences and Veterinary Public Health, Swedish University of Agricultural Sciences, Box 7028, 750 07 Uppsala, Sweden; SLU Global Bioinformatics Centre, Department of Animal Breeding and Genetics, Swedish University of Agricultural Sciences, Box 7023, 750 07 Uppsala, Sweden; Consejo Nacional de Investigaciones Científicas y Técnicas, Godoy Cruz 2290, 1425 Ciudad Autónoma de Buenos Aires, Argentina; Instituto de Agrobiotecnología y Biología Molecular, IABiMo, INTA-CONICET, Calle Las Cabañas y Los Reseros s/n, Casilla de Correo 25, Castelar, 1712, Buenos Aires, Argentina

**Author notes:** All authors contributed equally to this work.

## Abstract

Host associated microbiome not only may affect the individual health-status or provide insights into the species- or group specific bacterial communities but may act as early warning signs in the assessment of zoonotic reservoirs, offering clues to predict, prevent and control possible episodes of emerging zoonoses. Bats may be carriers and reservoirs of multiple pathogens such as viruses, bacteria and parasites, showing in the same time robust immunity against many of them. The microbiota plays a fundamental role on the induction, training and function of the host immune system and the immune system has largely evolved in order to maintain the symbiotic relationship of the host with these diverse microbes. Thus, expanding our knowledge on bat-associated microbiome it can be usefully in understanding bats’ outstanding immune capacities. The aim of this study was to investigate the presence of different bacterial communities in heart tissue of insectivorous bats, *Nyctalus noctula, Pipistrellus pipistrellus* and *Rhinoplophus hipposideros*, from Central and Eastern Europe using high-throughput sequencing of variable regions of the 16S rRNA. In addition, species-specific PCRs were used to validate the presence of the vector-borne pathogens *Bartonella* spp. and *Rickettsia* spp. In this study we identified a wide variety of bacterial groups, with the most abundant phyla being Proteobacteria and Firmicutes. The results showed that at individual level, the year or location had no effect on the diversity and composition of the microbiome, however host species determined both structure and abundance of the bacterial community. We report the presence of vector-borne bacteria *Bartonella* spp. in samples of *N. noctula* and indications of *Rickettsia* spp. in *R. hipposideros*. Our results provide a first insight into the bacterial community found in heart tissue of bats from Central and South Eastern Europe.

## Introduction

Bats have a series of unique characteristics among mammals. They show remarkable levels of tolerance to most known pathogens, high mobility (active flight), increased longevity compared with other small mammals and reduced senescence (compared to their size, Foley et al. 2019). Bats are the source for many diseases, like for example viruses causing rabies, EBOLA, severe acute respiratory syndrome, and Middle East respiratory syndrome all likely originating in bats (Hayman 2019). Moreover, bats are reservoirs of highly diverse pathogenic bacterial groups, without showing any apparent signs of disease (*Bartonella* spp., Corduneanu et al. 2018 or haemoplasmas - Holz et al. 2019). Bats are able to sustain an energetically demanding lifestyle attained through high basal metabolic rate (active flight). The most non-tropical bats are able to reduce their metabolic rate to virtually non-existent (while hibernation) in a short time (Rogers et al. 2019). Thus, a microbiome adapted to such an organism is interesting to study. Moreover, European bats are carriers or reservoirs for multiple zoonotic bacterial pathogens like *Bartonella* spp., *Leptospira* spp., *Rickettsia* spp. (Mühldorfer et al. 2011, Veikkolainen et al. 2014, Lilley t al. 2017) and the survey of these animals and their associated pathobiome may help to better understand their role in possible emerging zoonoses.

In bats, most high-throughput sequencing studies were focused on the identification and characterization of viruses. These studies used samples of saliva, urine (Fischer et al. 2015), guano (Li et al. 2010, Ge et al. 2012, Dufkova et al. 2015, Fischer et al. 2015), rectal swabs (Tse et al. 2012) or internal organ tissues (Dacheux et al. 2014) from different parts of the world, with a main focus on bats in the Yangochiroptera suborder (He et al. 2013, Dacheux et al. 2014, Donaldson et al. 2010, Li et al. 2010). On the other hand, studies on bacterial pathogens in bats have been primarily performed using species-specific PCR-based detection tests (Concannon et al. 2005, Kamani et al. 2014, Cicuttin et al. 2017), while metagenomics may be used as a tool to discover new bacteria and viruses that may affect animals and may be transmitted to humans (Blomström et al. 2010, Blomström et al. 2011, Belák et al. 2013, Franco Filho et al., 2019). Studies targeting the bacteriome of bats are a handful and these used samples of excreta (De Mandal et al. 2015, Dietrich et al. 2015), skin and fur (Avena et al. 2016, Lemieux-Labonté et al. 2016, Winter et al. 2017) or ectoparasites (Wilkinson et al. 2016).

To our knowledge this is the first investigation of the bacterial community of bat organs, and also the first bacteriome study conducted in European bats. The aim of our study was to (i) analyze the diversity and composition of the bacterial communities in bat heart tissue, (ii) to assess differences between different individuals (intraspecific) and species (interspecific), (iii) to test the metagenomics approach to validate the presence of vector-borne pathogens in tissues of bats.

## Materials and methods

### Source of tissue samples

We used tissues of bat corpses collected from Czech Republic and Romania in the period 2012-2013 and stored deep-frozen (−20 °C). All animals were found dead due to natural causes (sudden variation in temperature in the early emerging period) and collected below urban roosts (near buildings - Czech Republic) or close to the entrance of natural roosts (rock-crevices, in case of Cheile Bicazului and Huda lui Papara, both important hibernating sites, regularly visited by bat researchers in wintering and early emergence periods - Romania). Most bats were collected on the day of their death, while in case of Cheile Bicazului we estimate that some bat corpses were older than one day. However, in this case emerging bats fell into deep snow and were collected in frozen state on 30 March 2012. Here three sunny days of unusually high temperature were registered in late March 2012 (with midday temperature of 16 °C on 26^th^ and 27^th^, while 30 cm deep snow persisted in the gorge) increasing the temperature of hibernating crevices located high in limestone cliffs, which drove bats to initiate feeding forays. This was followed by a sudden drop in temperature on the night from 28^th^ to 29^th^ with the midday temperature on 30^th^ of March being 0 °C and below freezing point in the nights. While most bats managed to re-enter hibernation, we found many corpses (>300) and several weak individuals still clinging to the rocks below the roosts on the days following the warm-spell. Samples were collected from a selected number of organs (heart, liver, spleen and kidney) using scalpels. For each animal 2 sterile scalpel blades were used: first to cut the fur and skin, while the second to harvest the organ samples. All samples were collected in individual sterile zip bags and stored at −20°C, immediately after the necropsy. Bats were identified to species level using morphological keys (Dietz and von Helversen 2009). No ethical permit was necessary to perform corpse collection or necropsy in these countries.

### DNA extraction, PCR amplification

Genomic DNA was extracted from 25 mg of heart tissue using DNeasy Blood & Tissue Kit (Qiagen, Hilden, Germany) according to the manufacturer’s instruction and stored at −20 °C. For the 16S metagenomics studies, 31 bats out of 106 were randomly chosen, and they were from Czech Republic (n=1, Ochoz: 49.6001 N; 16.9143 E) and Romania (n=30; Cheile Bicazului: 46.8121 N; 25.8257 E and Huda lui Papară: 46.3814 N; 23.4616 E).

For these samples, an initial screening was performed using a specific PCR targeting the V3-V4 region of the 16S rRNA gene in accordance with the Illumina *16S Metagenomics Sequencing Library Preparation* guide (www.support.illumina.com). Amplification products were visualized by gel electrophoresis on 2% agarose gel stained with GelRed^®^ Nucleic Acid Gel Stain (Biotium Inc., Fremont, CA, USA).

From the 31 samples tested, 9 had an intense positive band, 22 had a weak positive band. From the samples that had weak band 3 pools were made, according to the species and the locations from where the samples were collected (Table 1). To concentrate the DNA of the weak band samples, DNA precipitation was performed. The protocol used for DNA precipitation included the following steps: for each sample a 1/10 volume of sodium - acetate (3M) was added (pH=5.2), followed by the adding of 2.5 volume of ice-cold 100% ethanol. Each sample was mixed and stored at −20 °C for at least one hour to precipitate the DNA. The precipitated DNA was recovered by centrifugation at full speed in a cold microcentrifuge for 15 minutes. The ethanol was then removed, and the pellet was washed twice with 70 - 75% ethanol at room temperature. The DNA pellet was allowed to air-dry and then suspended in 25 µl of sterile TE buffer.

**Table 1.** Species, sampling location and country of samples sequenced using Ion S5XL platform.

| ID/ Type of sample | Bat species | Location | Country |
| --- | --- | --- | --- |
| 1/ Individual | *Nyctalus noctula* | Cheile Bicazului | Romania |
| 2/ Individual | *Nyctalus noctula* | Cheile Bicazului | Romania |
| 3/ Individual | *Nyctalus noctula* | Cheile Bicazului | Romania |
| 4/ Pool | *Nyctalus noctula* | Cheile Bicazului | Romania |
| 5/ Individual | *Pipistrellus pipistrellus* | Huda lui Papară | Romania |
| 6/ Individual | *Pipistrellus pipistrellus* | Huda lui Papară | Romania |
| 7/ Individual | *Pipistrellus pipistrellus* | Huda lui Papară | Romania |
| 8/ Individual | *Pipistrellus pipistrellus* | Huda lui Papară | Romania |
| 9/ Pool | *Pipistrellus pipistrellus* | Huda lui Papară | Romania |
| 10/ Individual | *Rhinolophus hipposideros* | Ochoz | Czech Republic |

### DNA library construction and sequencing

The selected 12 samples (9 individual and 3 pools with 7-8 samples/pool) were PCR amplified using the multi-primer bacterial Ion 16S™ Metagenomics Kit, following the kit protocols for library preparation. The kit includes 2 sets of primers: V2, V4, V8 and V3, V6-7, V9, corresponding to the hypervariable regions of the 16S rRNA gene in bacteria. The quality assurance and high-throughput sequencing was performed by Uppsala Genome Centre (UGC), which is a part of the Science for Life Laboratory (SciLifeLab, Uppsala, Sweden) using the Ion S5XL platform with an Ion S5 530 Chip and 400bp read length chemistry. The quality of the sequencing process was analyzed using Ion Reporter (v5.0) metagenomics 16S workflow (ThermoFisher Scientific, Waltham, MA, USA, Table 2). Two samples (one individual and one pool) failed, thus resulting 10 sequences in total (Table 1).

**Table 2.** Run report results for the sequencing quality using Ion S5XL.

| Sample ID | Bases | >= Q20 | Reads | Mean Read length |
| --- | --- | --- | --- | --- |
| 1/ Individual | 258,598,494 | 237,480,576 | 1,125,592 | 230 bp |
| 2/ Individual | 183,381,574 | 165,323,553 | 853,770 | 215 bp |
| 3/ Individual | 311,182,533 | 284,710,797 | 1,324,145 | 235 bp |
| 4/ Pool | 221,179,881 | 203,492,857 | 959,196 | 231 bp |
| 5/ Individual | 243,697,521 | 223,031,079 | 1,083,876 | 225 bp |
| 6/ Individual | 221,470,365 | 201,973,762 | 976,028 | 227 bp |
| 7/ Individual | 252,728,506 | 229,928,240 | 1,110,718 | 228 bp |
| 8/ Individual | 225,325,799 | 203,046,147 | 962,043 | 234 bp |
| 9/ Pool | 332,469,317 | 303,815,851 | 1,453,701 | 229 bp |
| 10/ Individual | 87,177,417 | 78,486,446 | 416,313 | 209 bp |

### Microbial Community Analysis

Microbial composition and diversity of 7 variable regions were analyzed only for samples that were successfully amplified and sequenced (ID 1-10, Table 1), using Quantitative Insights into Microbial Ecology (QIIME) v1.9.1 (Kuczynski et al. 2012) following the pipeline described by Barb *et al.* (2016) and the Brazilian Microbiome Project (BMP) (Pylro et al. 2014). Briefly, the sequences were pre-processed for quality filtered using Trimmomatic v0.33 (Bolger *et al.*, 2014), trimming if the average quality within a 4-base window falls below Q20 and removing reads shorter than 200 bp.

The sequences were concatenated into one *fasta* file containing reads from all samples and all different variable regions. While the 16S Metagenomics kit contains 7 different variable regions, the same primer pair targets V6 and V7, thus we will refer to 6 regions. These regions were split aligning all reads with Mothur script *align.seqs* using Silva as the reference database (www.mothur.org/, Silva version 128) with the default parameters. The forward and reverse reads were grouped into their corresponding regions based on where they aligned using the 16S rRNA gene coordinates following Barb *et al.* (2016).

The data analysis workflow was run 6 separate times in order to create operational taxonomic unit (OTU) tables for each region. Chimera checking due to PCR artifacts was performed using the UCHIME algorithm and then the open-reference OTUs picking was performed using UCLUST algorithm. Taxonomy was assigned against the Silva reference OTU build version 128, using a 97% sequence similarity threshold. Singletons and OTUs with abundance below 0.005% were filtered out from the final OTU tables (Edgar 2013).

The workflow was continued in QIIME performing the diversity analysis, comparing within-samples (alpha diversity) through richness (Chao 1 index and observed OTUs) and community diversity (Shannon and Simpson indexes) for each region. Furthermore, principal coordinate analysis (PCoA) plots were generated in QIIME based on weighted and unweighted UniFrac distance matrix for all regions. This method is a β-diversity measure that takes into account the phylogenic divergence between OTUs to identify differences in the overall microbial community structure between samples (Lozupone et al., 2011).

### PCR for detection of bacterial pathogens

A number of conventional PCRs (cPCR) targeting the *glt* A, *rpo* B and ITS encoding genes of *Bartonella* spp. were used for validation. For *Rickettsia* spp. two cPCR-s targeting the *glt* A and *omp*A genes were used (all primers, with summary protocols are listed in Supplementary file 1). The reactions were carried out in with 25 µl reaction mixture which contained 2x Green Master Mix (Rovalab GmBH, Teltow, Germany), 6.5 µl water, 0.01 mM of each primer, 4 µl of DNA. The PCR was performed using the T^1000TM^ Thermal Cycler (Bio-Rad, Hercules, CA, USA) with the conditions specific for each set of primers (Supplementary file 1). The positive controls used were *B. henselae* isolated from a cat and *R. helvetica* isolated from a tick (*Ixodes ricinus*). For each reaction two negative controls were used. Amplification products were visualized by electrophoresis on 1.5% agarose gel stained with RedSafe™ 20,000× Nucleic Acid Staining Solution (Chembio, St Albans, UK), and their molecular weight was assessed by comparison to a molecular marker (Hyperladder IV, Bioline, London, UK).

PCR products were purified using a commercial kit (Isolate II PCR and Gel Kit, Bioline, London, UK) and sent for sequencing (Macrogen Europe, Amsterdam, The Netherlands). The sequences were compared with those available in GenBank^®^ using Basic Local Alignments Tool (BLASTn) analyses.

## Results

In total, eight individual and two pooled samples from three different bat species were analyzed using multi-primer bacterial Ion 16S™ Metagenomics (Table 1). All sequence data were deposited in the NCBI Sequence Read Archive database under the BioProject accession number PRJNA623883. The accession numbers for the raw sequences are the following: SAMN14560019-SAMN14560028.

In these samples, a great intraspecific variation was found (Table 3). Proportional abundance within each sample for each region was determined for phyla and class levels and stacked bar charts at both levels are shown (Fig. 1 and 2). The minimum and maximum relative abundance values for the different variable regions sequenced are shown in Table 3. The most abundant phyla were Proteobacteria, Firmicutes and Actinobacteria (Fig. 1), with predominance of the Gammaproteobacteria, Bacilli and Actinobacteria classes respectively (Fig. 2) in all variable regions of 16S rRNA studied. Analyzing only the V4 region by host species, the highest relative abundance of bacterial phyla was also represented by Proteobacteria, (class Gammaproteobacteria, Fig. 3 and 4).

**Table 3.**
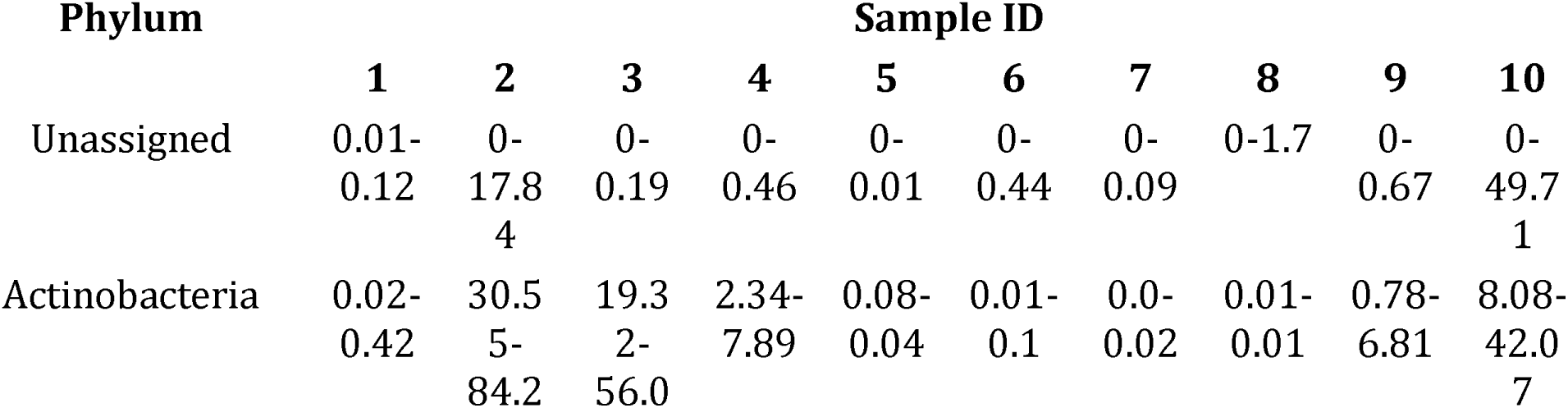
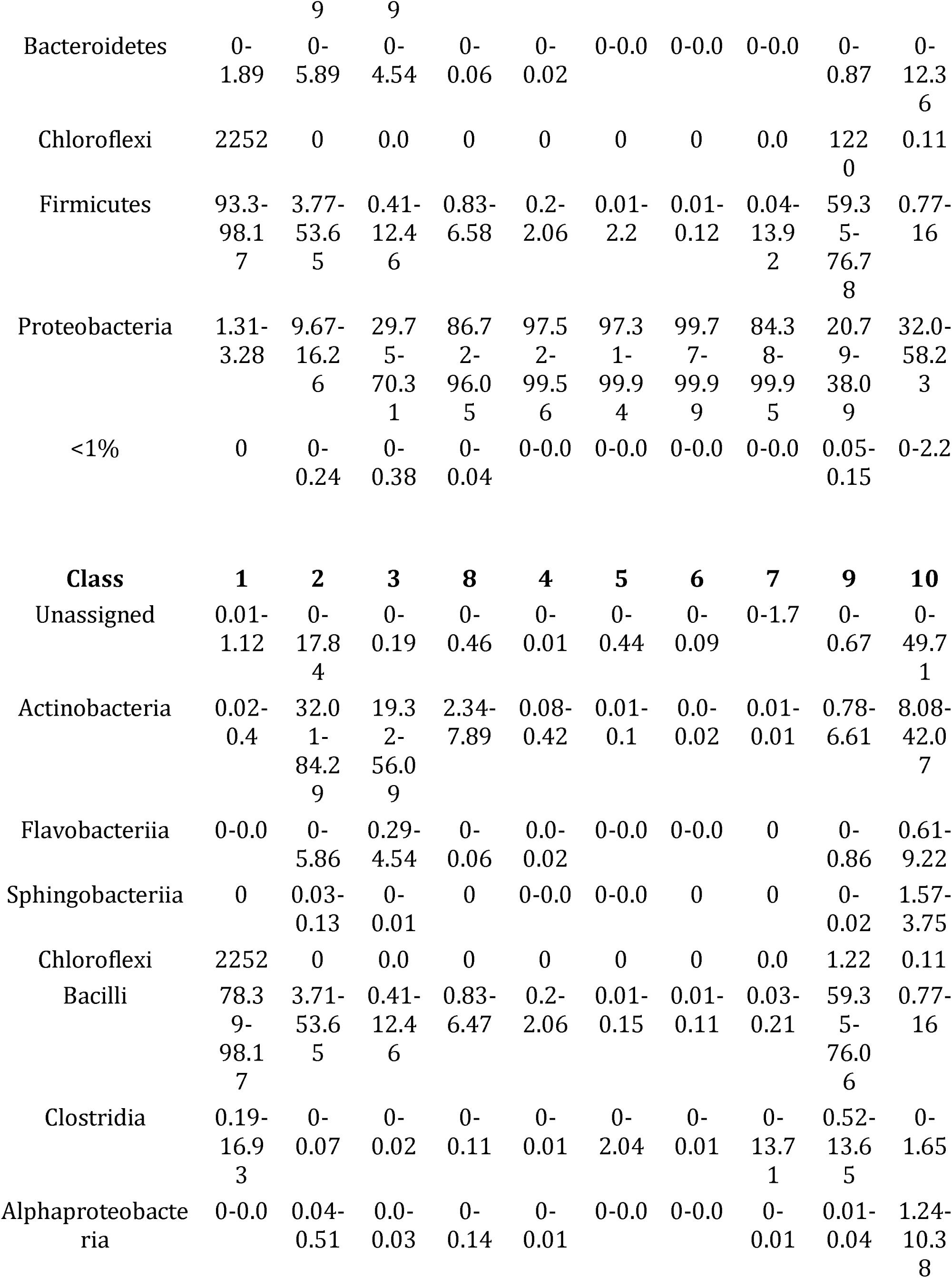
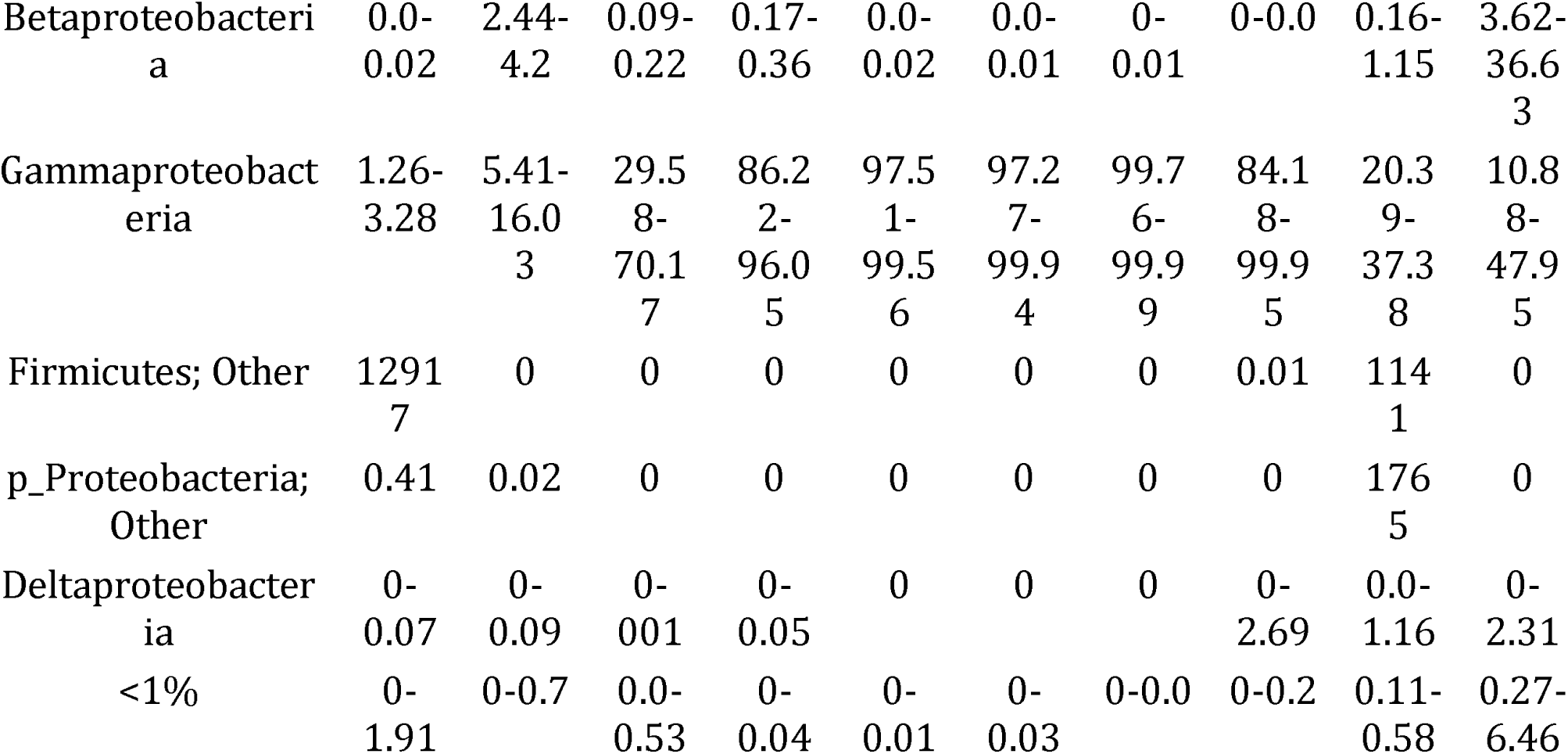
Phylogenetic structuring of important bacteria detected, with the minimum and maximum relative abundance recorded across all regions at sample level

**Fig 1.**
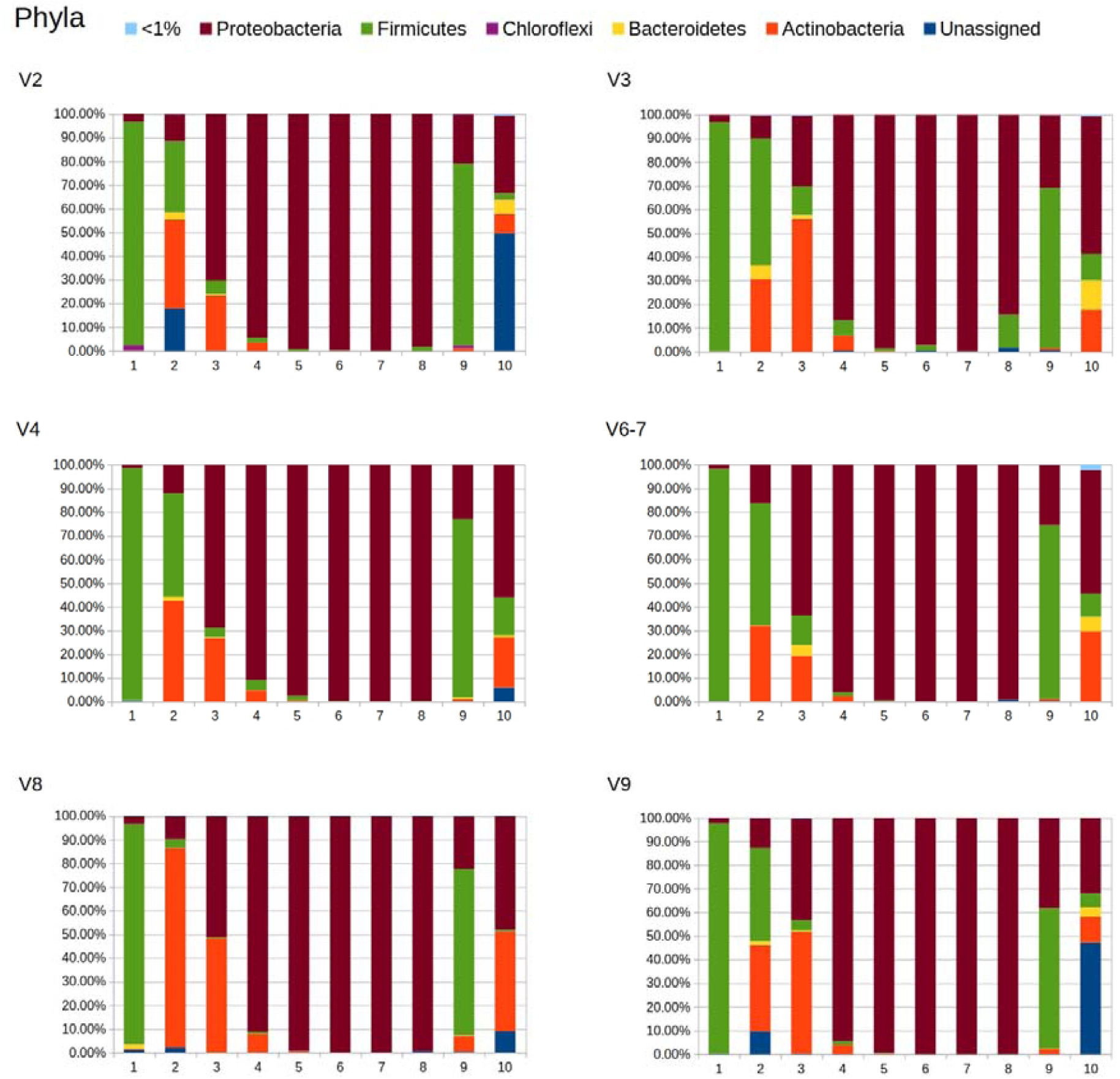
Relative abundance of bacterial phyla in bat heart samples (all variable regions).

**Fig 2.**
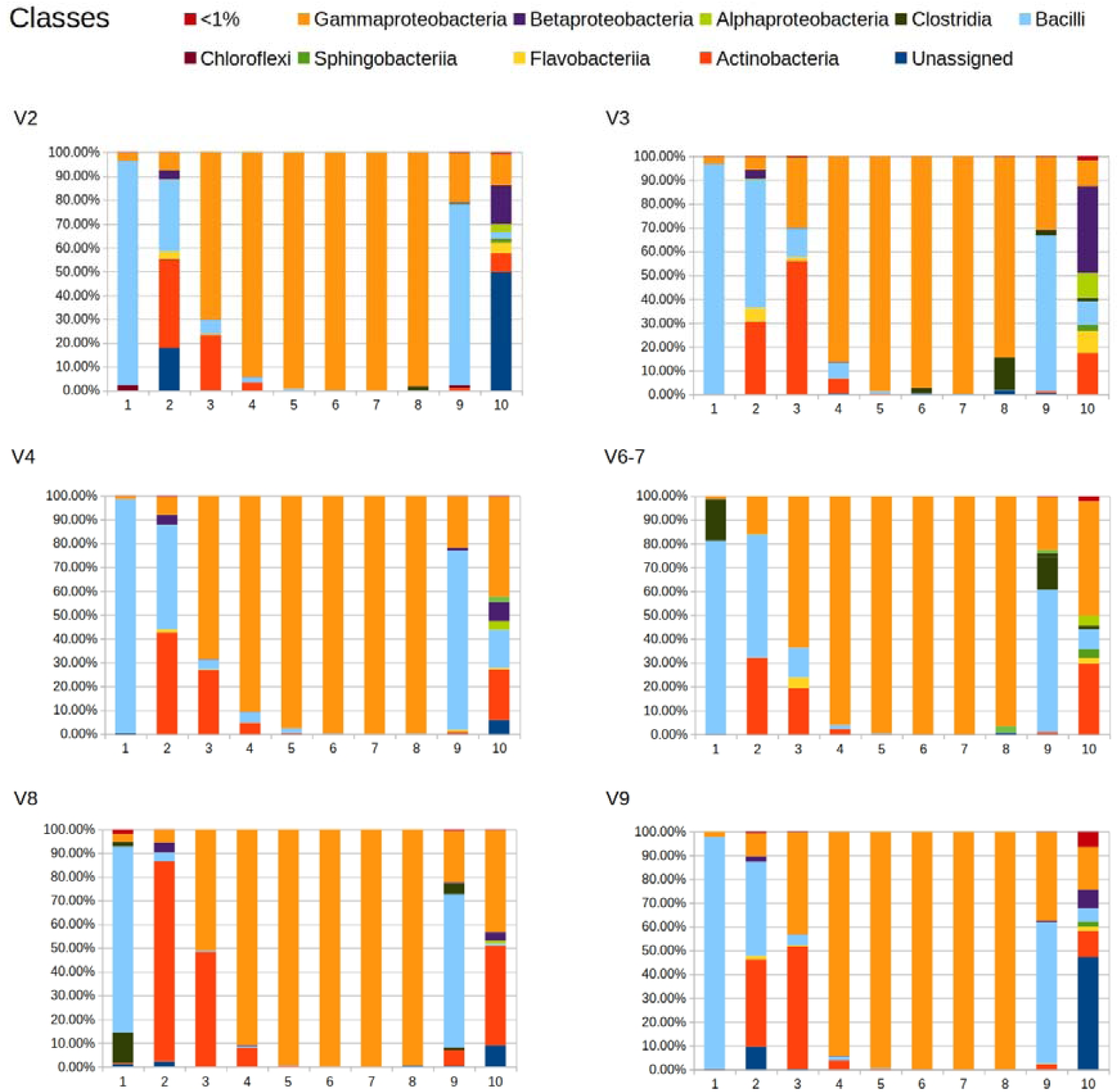
Relative abundance of bacterial classes in bat heart samples (all variable regions).

**Fig 3.**
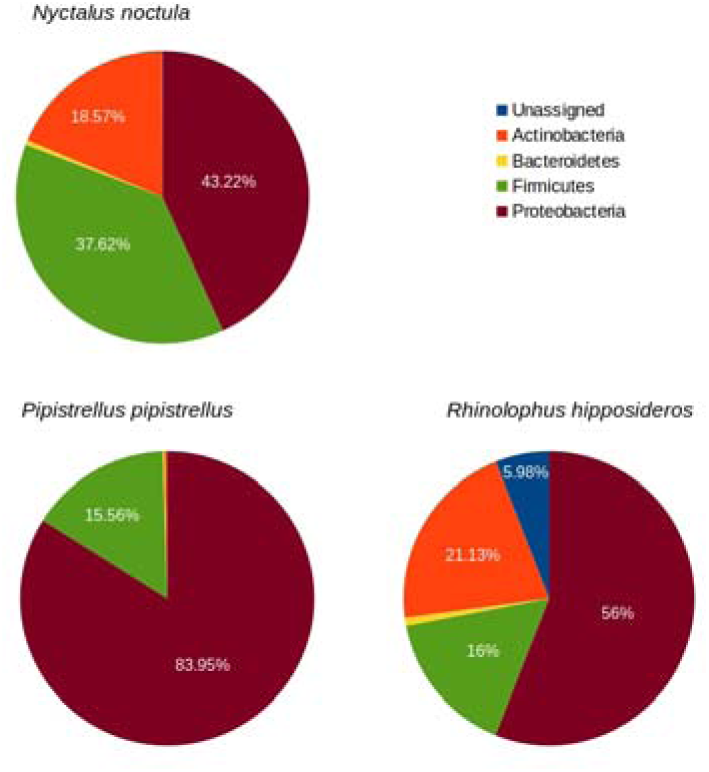
Relative abundance of the five dominant bacterial phyla targeting the V4 region in bat heart tissues.

**Fig 4.**
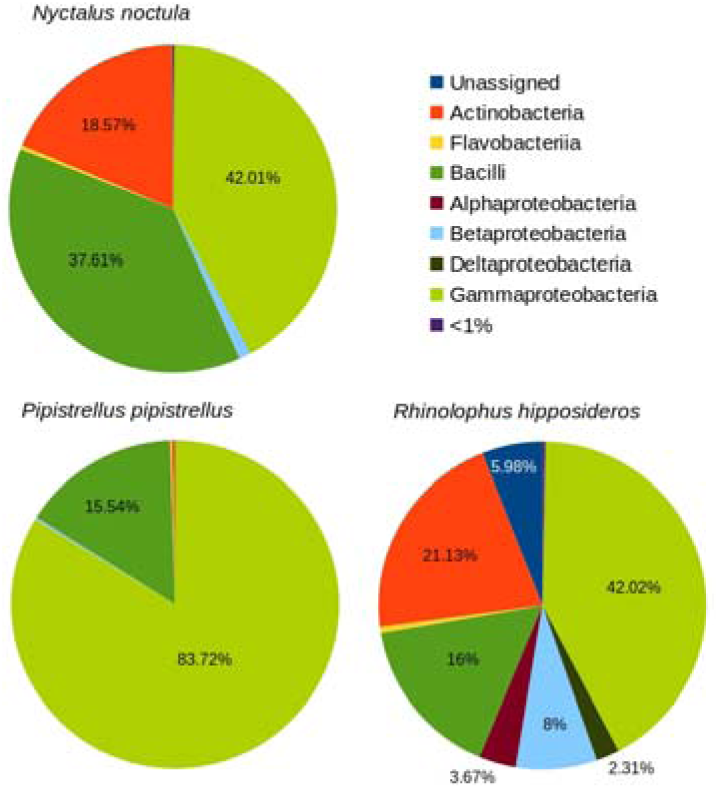
Relative abundance of the nine dominant bacterial classes targeting the V4 region in bat heart tissues.

In the samples of *P. pipistrellus* only Proteobacteria and Firmicutes were detected, represented by Gammaproteobacteria and Bacilli classes, while Actinobacteria appeared as one of the predominant phyla in *N. noctula*, along with Proteobacteria and Firmicutes (Fig. 4 and 6). In *R. hipposideros* the bacteriome showed the widest diversity at phyla level, dominated by the Proteobacteria (Fig. 5). This was also evident in alpha diversity indexes, which showed no patterns related to locality or year, but were determined by host species (observing the highest values of Shannon and Simpson diversity index for *R. hipposideros*, Table 3 and on-line Supplementary material 2, Fig 7).

**Fig 5.**
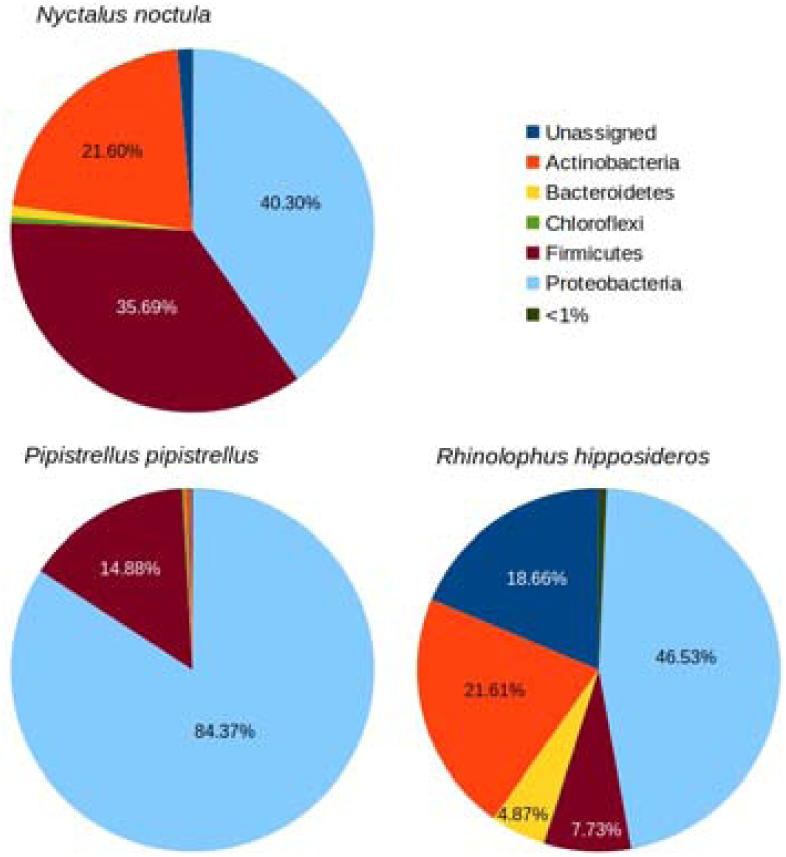
Relative abundance of the five dominant bacterial phyla in bat heart tissues as the average of the six variable regions sequenced.

**Fig 6.**
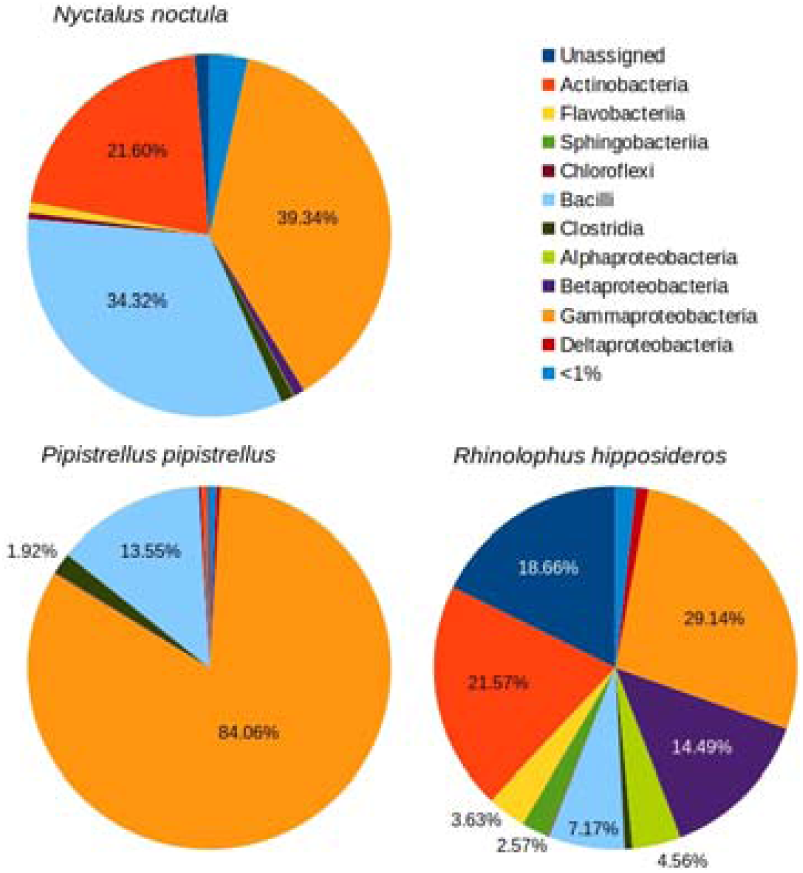
Relative abundance of the ten dominant bacterial classes in the three bat species (as the average of the six variable regions).

**Fig 7.**
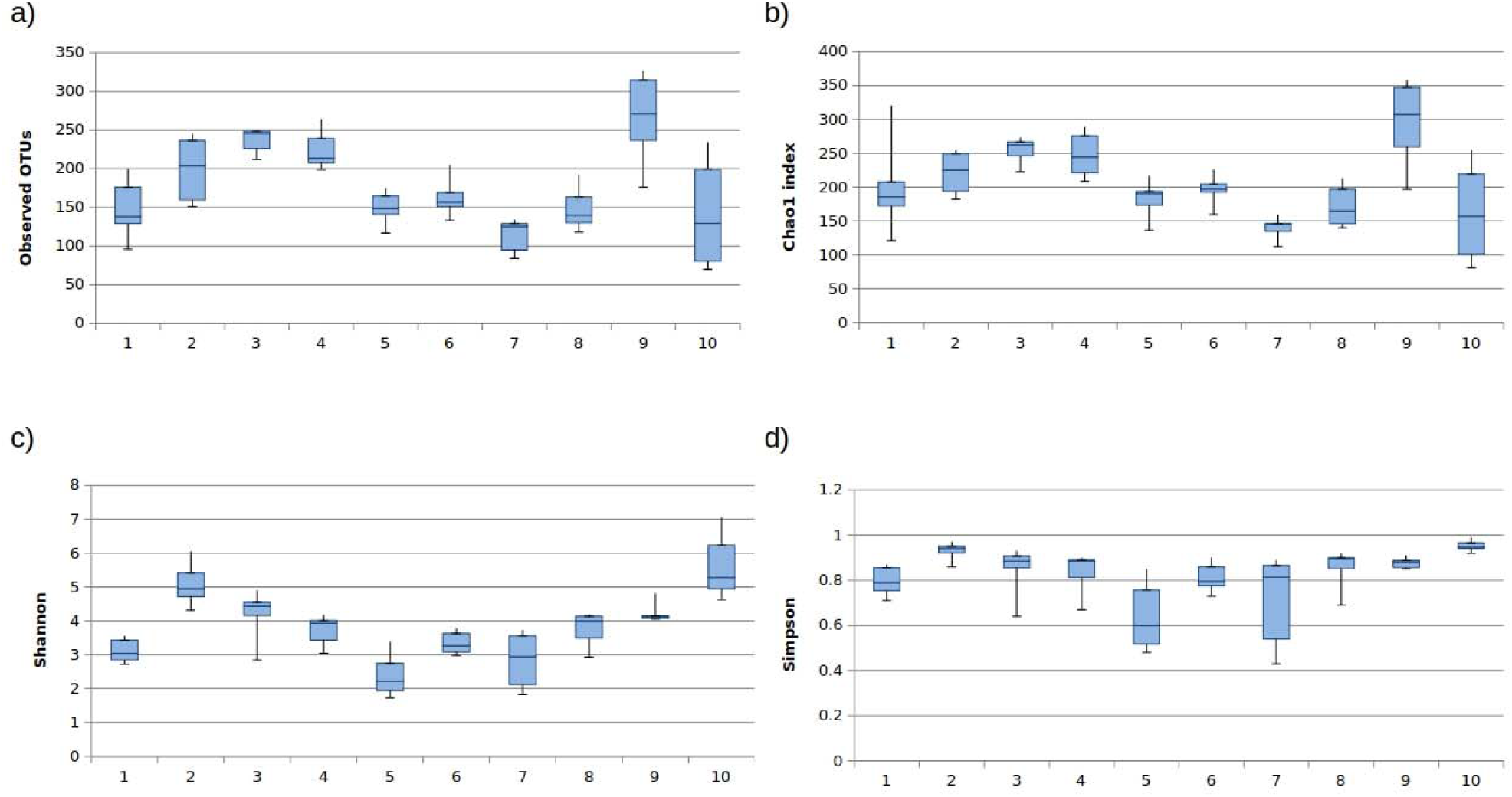
Alpha diversity of bacteria in hart tissues of bat samples calculated through richness (a. Chao1 index, b. observed OTUs) and community diversity (c. Shannon index, d. Simpson index)

A principal coordinate analysis (PCoA) based on weighted and unweighted UniFrac distances were conducted for each region to determine any separation into sample clusters. Only the PCoA plots for the V4 variable region revealed that the samples corresponding to each species (Sp) or locality differentially modulate the microbiota of bats (Fig 8). The ADONIS analysis detected significant changes in β-diversity only in the unweighted UniFrac among species (p = 0.05894) and Locality variables (p = 0.08192), which is consistent with the structure of the data depicted in the PCoA plot.

**Fig 8.**
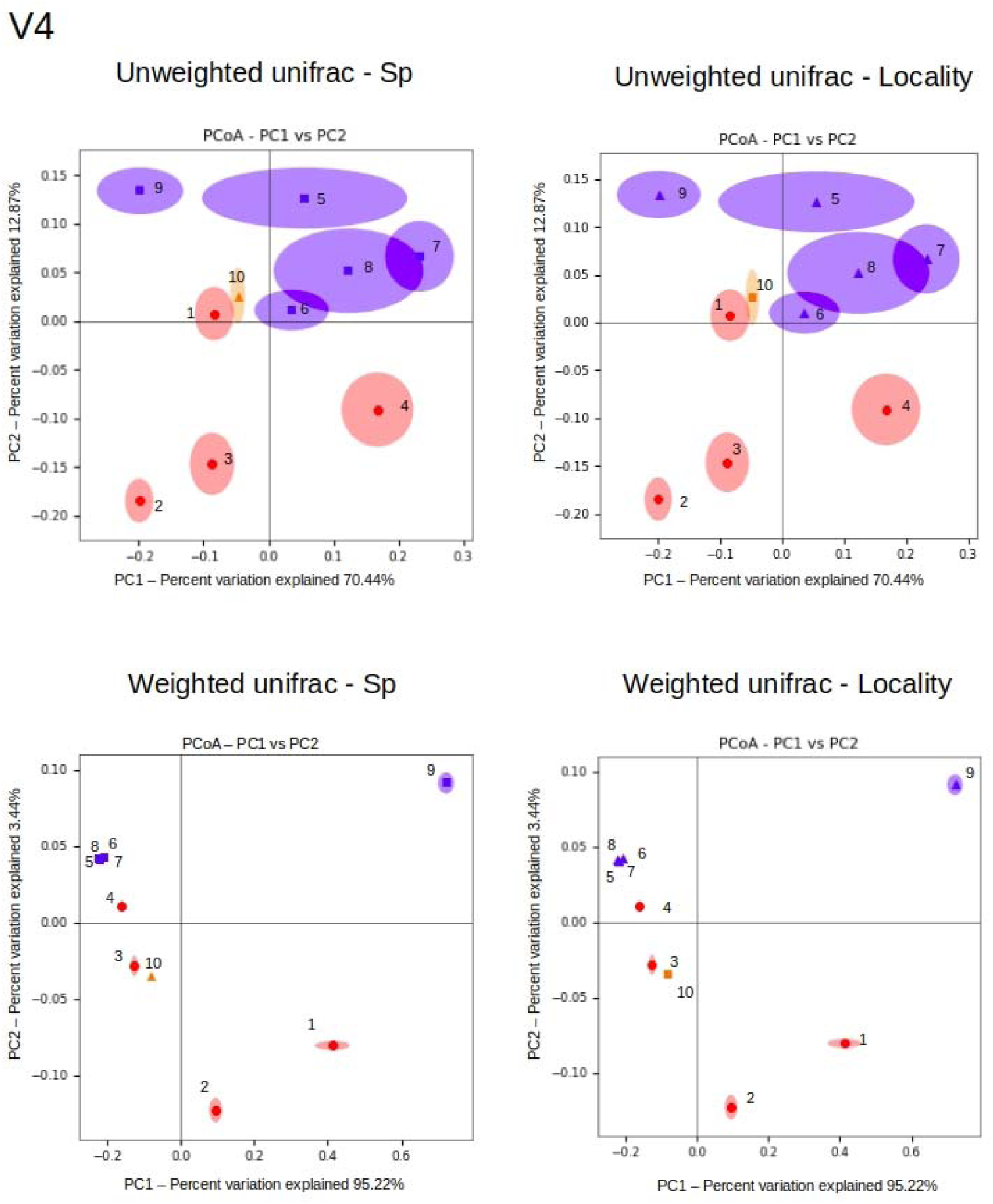
Principal coordinates analysis (PCoA) of weighted and unweighted UniFrac distances using V4 variable region.

Most identified bacterial genera host primarily commensals and saprophyte bacteria; however several pathogenic or parasitic genera were recorded (Table 4). Using targeted PCR two intracellular pathogens *Bartonella* spp. (pooled samples 4 – *N. noctula* and 9 – *P. pipistrellus*) and *Rickettsia* spp. (sample 10 – *R. hipposideros*) were recorded, but in very low proportions (<0.005% relative abundance). These results were subsequently evaluated using the primers shown in Supplemental material 1. Each individual sample from the pools 8 and 9 was tested individually for the presence of *Bartonella* spp. targeting different genes and 2 samples were positive (one targeting the *glt*A gene and one the ITS gene). Unfortunately, it was not possible to amplify *Rickettsia* spp. from the sample 10. After Sanger sequencing only one sample was positive for the presence of *Bartonella* spp. targeting the ITS gene (Accession Number: MK828122). BLASTn analysis of the ITS sequence showed to be most similar (93.27% identity) to the uncultured *Bartonella* sp. (GenBank^®^, KX420720.1) from *Myotis emarginatus* in Georgia, Caucasus.

**Table 4.**
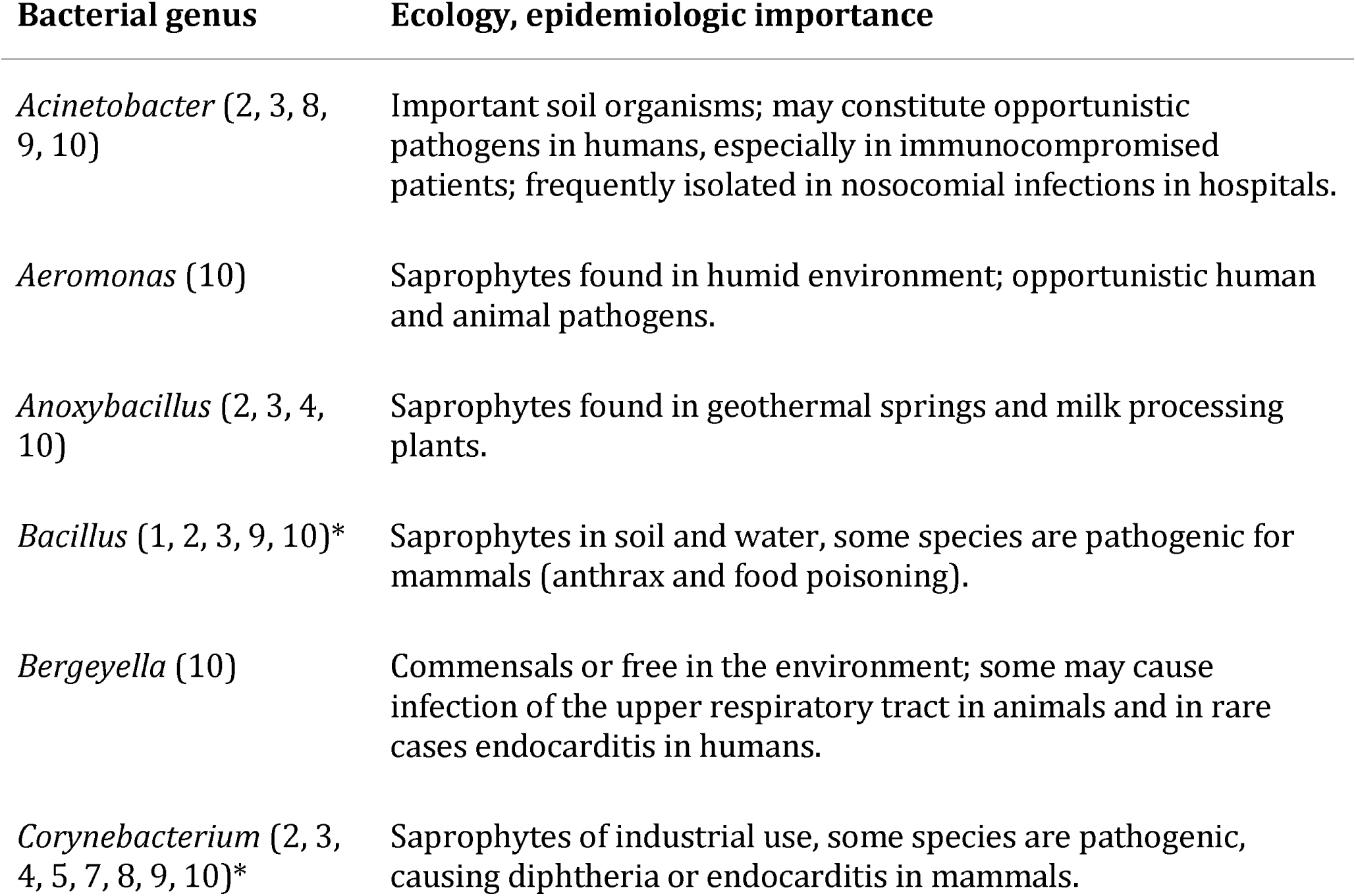
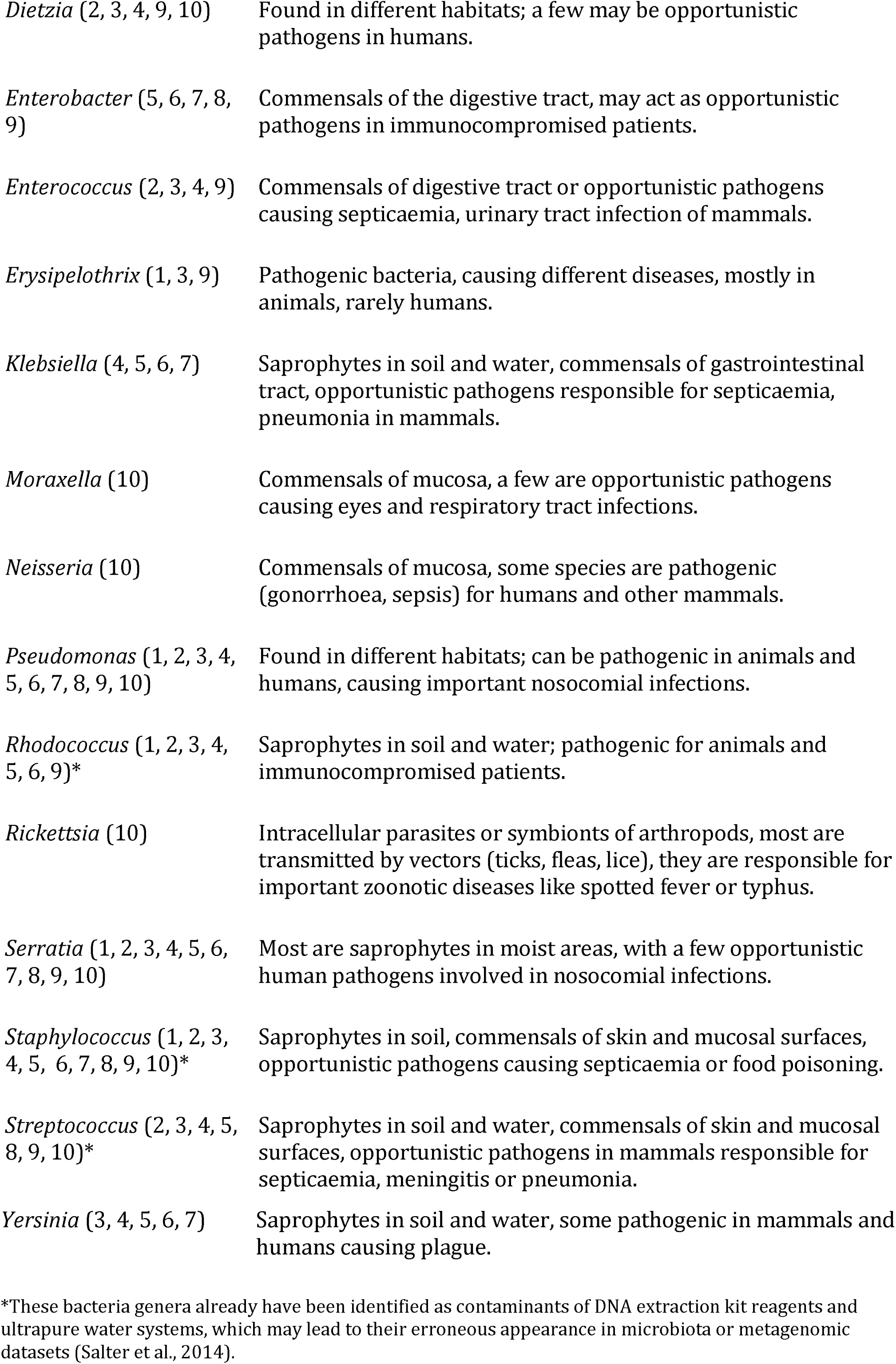
The most common bacterial genera detected, with sample ID of dominant presence and ecological and epidemiological importance.

## Discussion

We studied bacterial communities present in heart tissues of insectivorous bats from Central and South Eastern Europe, with a focus on three bat species (*N. noctula, P. pipistrellus* and *R. hipposideros*). Little is known about the ecology of bats’ associated bacteria and the role that these play in bats’ health or their possible role as pathogenic reservoirs. By assessing the bat-bacteriome one may shed light not only on the commensals of the hosts, but also on the role these animals may play in the maintenance (and possible shedding) of pathogenic bacteria (He et al. 2013, Wilkinson et al. 2016). Our samples came from common European bat species (Dietz and von Helversen 2009), and even from bats sampled in urban areas.

Metcalfe et al. (2016) showed that microbial community undergoes a continual change along the process of carcass decomposition, their results showed the first changes occurring after the 4^th^ day of carcass decomposition as in lab, as in field conditions. In our case most carcasses were collected within one day of decease, so we are fairly confident that our results show a bacterial diversity which is close to the bacteriome of live bats.

Previous studies on the bacterial communities present in different samples collected from bats targeted different regions of the 16S rRNA gene (maximum two), using different sequencing technologies (pyrosequencing, sequencing by synthesis). Most studies were performed using Illumina next generation sequencing (targeting the v3-v4 region) (De Mandal et al. 2015, Avena et al. 2016, Lemieux-Labonté et al. 2016, Dietrich et al. 2017, Dietrich and Markotter 2019). We chose multi-primer bacterial 16S™ Metagenomics for this study because it targets altogether 7 different hypervariable regions of the 16S rRNA gene (in comparison to previous studies where one or two regions were targeted, such as V1, V2 (Winter et al. 2017), V3 and/ or V4 region (Lemieux-Labonté et al. 2016, Wilkinson et al. 2016). This approach allowed us to get a better overview of the diversity of bacterial communities present (Barb et al. 2016). Examples of 16S rRNA studies in bats using the V4 region were the assessment of the bats’ skin microbiome in North America (Avena et al. 2016, Lemieux-Labonté et al. 2016) and the microbiome present in guano collected from a cave in Meghalaya, India (De Mandal et al. 2015). Also, the same V4 region was studied to determine the gut microbiota and the associated diet of several bat species in the Neotropics (Carrillo-Araujo et al. 2015). In other studies, two regions were assess, such as V3 and V4 to determine the bacteria present in bat flies from Madagascar (Wilkinson et al. 2016) and from other biological samples (saliva, urine, feces, see Dietrich et al. 2017), while the V1-V2 region was targeted to determine the skin and fur microbiome of bats from North America (Winter et al. 2017).

There are multiple factors influencing the community structure when the bacteriome of a sample is analyzed, such as DNA extraction method, different regions of the 16S rRNA targeted, primers used, the type of sequencer (Illumina, IonTorrent, 454) and the bioinformatics pipelines and parameters used (Kim et al. 2011, Kennedy et al. 2014, Fouhy et al. 2016). For this study we used the heart tissue, which contains a relative high amount of blood, thus increasing the chances of detecting vector-borne bacteria. We compared our results with other studies using all regions of the 16S rRNA gene sequenced, and an extraction of only V4 region information.

The bacterial phyla present in our samples were Proteobacteria, Firmicutes (the two most abundant), Actinobacteria, Bacteroides and some belonging to unassigned classes. The situation was similar when we analyzed only the V4 region, showing that Proteobacteria and Firmicutes had the highest bacterial prevalence (Fig. 7). The phylum Proteobacteria is frequently found to be dominant in bats, like in saliva of bats from South Africa (>90%) (Dietrich et al. 2017). Our study showed that Gammaproteobacteria was the most abundant class from the phyla Proteobacteria in *P. pipistrellus* samples, either if we analyzed all regions or only the V4 region (84.06% and 83.72% respectively) and Bacilli showed the highest proportion in the heart tissue of *N. noctula* (34.32% and 37.61%) (Fig. 6, 8). The sample belonging to *R. hipposideros* showed the highest diversity of dominant bacterial classes (Fig. 5, 7) and this may correlate with specific ectoparasite diversity/abundance, as ectoparasites were shown to enhance the bacterial diversity in the host’s blood (McKee et al. 2019). The most abundant phylum of the skin microbiome of bats and humans is represented by Actinobacteria (Lemieux-Labonté et al. 2016, Winter et al. 2017, Oh et al. 2012). We observed Firmicutes in all bat species, with the highest value in *N. noctula* (35.69% for all regions and 37.62% for V4) (Fig. 5 and 7). This was the most abundant phylum in fecal and urine samples of bats, too (Dietrich et al. 2017).

We noted high presence of several classes of decomposing bacteria, such as those from the phylum Proteobacteria (specifically, *Pseudomonas*) or from the phylum Firmicutes (eg. *Clostridium*) (Hyde et al. 2013). While most species of both mentioned bacterial genera are decomposing, also a number of important animal (and human) pathogenic species belong to these genera. Thus, although we are not able to rule out the start of decaying of the bat’s internal organs (due to the time lag between the animal death and the collection), the high levels may result from earlier infections, which could have contributed to the bats decease (Mühldorfer et al. 2013).

Reports of *Bartonella* spp. infections are known from blood of bats (Judson et al. 2015, Han et al. 2017, Urushadze et al. 2017), from tissues (Bai et al. 2017, Stuckey et al. 2017) and from their associated ectoparasites (Morse et al. 2012, Davoust et al. 2016, Sándor et al. 2018, Hornok et al. 2019). In our study this pathogen group was found in low proportions, in two samples (2 out of 10, 20%), but the PCR targeting the ITS gene of *Bartonella* spp. was positive for one sample belonging to *N. noctula* from Romania. The study of microbiome from saliva, urine and feces of bats collected from South Africa showed a high percentage of this pathogen, ranging between 17% (saliva) and 67% (feces) (Dietrich et al. 2017). While our *Bartonella* spp. prevalence seem very low in comparison, insectivorous bats from Europe show differential levels of *Bartonella* spp. infection, with the here tested species showing the lowest prevalence even in live bat individuals (Corduneanu et al. 2018). The vector-borne pathogen *Rickettsia* could not be amplified by cPCR and maybe one reason is that we used primers for *Rickettsia* belonging to the spotted fever group, while in bat associated ectoparasites other *Rickettsia* spp. were found (Hornok et al. 2019). Identification of infectious bacteria from the bloodstream, particularly in humans are usually performed using blood cultures, but different studies compared this method with the 16S metagenomics, showing that this later technique is more sensitive and specific (Decuypere et al. 2016, Rutanga et al. 2018, Watanabe et al. 2018]. The blood tested for those studies was fresh however there are no studies that have tested the sensitivity in partly degraded heart or heart tissue.

There are multiple natural causes from which bats may die and testing tissues may provide clues for the presence of different pathogens. The greatest challenge is to quickly identify the dead animals to avoid decomposition of the body. Compared to the amount of research available on the human microbiome (Muszer et al. 2015), the microbiome of animals, and particularly of bats, remains relatively unknown, thus new information may be valuable and may provide us new insights into the understanding the complex relationship between the ecology of bacteria and their vertebrate host. Further studies should be conducted in order to identify the impact of microbiome on bats health and the role that can possibly have in transmission of different diseases.

## Conclusions

This study identified the bacterial communities in heart tissues of 3 species of insectivorous bats in Central and Eastern Europe, using 16S rRNA metagenomics approach. We found diverse bacterial communities, with the most abundant phyla being represented by Proteobacteria and Firmicutes. No effect regarding the bacterial community was observed at individual level, year or location, however host species determined both structure and abundance of the bacterial community. While most bacterial genera recorded may contain human pathogenic species, too, we report here only *Bartonella* spp. in samples of *Nyctalus noctula* and indications of possible presence of *Rickettsia* spp. in *Rhinolophus hipposideros*. These findings show one more time that bats are involved in the transmission of pathogens with a possible zoonotic impact.

## Acknowledgments

The authors would like to acknowledge the support of the National Genomics Infrastructure (NGI) / Uppsala Genome Center (funded by RFI/VR and Science for Life Laboratory, Sweden) and the SLU Bioinformatics Infrastructure (SLUBI) for providing assistance in massive parallel sequencing and computational infrastructure. Work performed at NGI / Uppsala Genome Center has been funded by RFI / VR and Science for Life Laboratory, Sweden. AC was supported by the COST Action TD1303: European Network for Neglected Vectors and Vector-Borne Infections (EURNEGVEC) and ERASMUS + program. The János Bolyai Research Scholarship of Hungarian Academy of Science, the ÚNKP 19-4-ÁTE-10 New National Excellence Program of the MIT and NKFIH 132794 provided financial resources to ADS, while SH received funding from NKFI 130216. MM received financial support from the Swedish Research Council for Environment, Agricultural Sciences and Spatial Planning, Formas (Grant Number 221-2012-586). NPV received funding from the Marie Curie International Research Staff Exchange Scheme within the 7th European Community Framework Program under grant agreement No PIRSES-GA-2013-612583-DEANN. The authors would like to thank for the kind contribution of Péter Estók, Sándor Boldogh (Hungary), Juliette Hayer and Tomas Klingstr⍰m (Sweden).

## Notes

### Competing Interest Statement

The authors have declared no competing interest.

